# Information transfer in mammalian glycan-based communication

**DOI:** 10.1101/2021.05.10.443458

**Authors:** Felix F. Fuchsberger, Dongyoon Kim, Natalia Baranova, Marten Kagelmacher, Robert Wawrzinek, Christoph Rademacher

**Affiliations:** Max-Planck-Institute of Colloids and Interfaces, Department of Biomolecular Systems, Am Mühlenberg 1 14424 Potsdam, Germany; Department for Pharmaceutical Chemistry, University of Vienna, Althanstrasse 14, 1090 Vienna, Austria

**Keywords:** C-type lectins, signal transduction, signal integration, glycobiology, innate immunity, signalling

## Abstract

Glycan-binding proteins, so-called lectins, are exposed on mammalian cell surfaces and decipher the information encoded within glycans translating it into biochemical signal transduction pathways in the cell. These glycan-lectin communication pathways are complex and difficult to analyse. However, quantitative data with single cell resolution provides means to disentangle the associated signalling cascades. We chose C-Type lectin receptors (CLRs) expressed on immune cells as a model system to study their capacity to transmit information encoded in glycans of incoming particles. Lectin receptor NFκB-reporter cell lines expressing DC-SIGN, MCL, dectin-1, dectin-2, and mincle, as well as TNFαR and TLR-1&2 in monocytic cell lines were characterized by comparing their efficiency to transmit glycan-encoded information. The information content was measured by following NFκB dependent GFP expression. While most receptors did transmit information to NFκB efficiently, we found dectin-2 to be an inefficient signalling receptor. Yet upon closer analysis we show that the sensitivity of the dectin-2 signal transduction pathway (EC_50_) can be enhanced by overexpression of its co-receptor FcRγ, while its transmitted information cannot. In this context, we expanded our investigation towards the integration of multiple signal transduction pathways, which is crucial during pathogen recognition. We show how lectin receptors using a similar signal transduction pathway (dectin-1 and dectin-2) are being integrated; by striking a compromise between the lectins. By using dectin-2 and other lectins as example we demonstrate how cellular heterogeneity and the receptor itself determine the efficiency and therefore outcome of the signal transduction pathways.

## Introduction

Glycans are present on all living cells and play a key role in many essential biological processes including development, differentiation and immunity. Being surface exposed, glycans often encode for information in cellular communication such as self-/non-self-discrimination, cellular identity and homing as well as apoptosis markers.(Bode et al., 2019; Maverakis et al., 2015; Williams, 2017) Other than linear biopolymers, such as proteins and nucleic acids, glycans are branched structures, where subtle changes in the glycosidic bonds between each monomer can carry essential pieces of information. Adding to this complexity, glycans are products of a large cellular machinery and are therefore not directly encoded by the genome.(Cummings, 2009) Besides their composition, the recognition of glycans by their receptors is complicated, particularly due to the lack of specificity: Glycans are recognized by lectins, yet no glycan is recognized by a single receptor and no individual lectin is highly specific for only one glycan. Additionally, affinities are low and interactions often depend on multivalency of both the receptor and the ligand. Overall, since alterations of the glycocalyx do not function as a deterministic on/off switch but rather a progressive tuning of the cellular response, glycan lectin communication should be considered as a stochastically behaving system, rather than a deterministic one.(Dennis, 2015)

Many lectin receptors serve as triggers for multiple immunological signalling pathways, often funnelling down to NFκB as a transcription factor. In this work we focus on C-Type lectin receptors (CLRs): Mincle for example is a CLR involved in the recognition of pathogens as well as self-damage.(Miyake et al., 2015; Williams, 2017) Mincle and its close relative dectin-2 signal *via* the FcRγ gamma chain (Miyake et al., 2015; Ostrop et al., 2015; Sato et al., 2006), leading to CARD9-BCL-10-Malt1 activation. This in turn results in activation of NFκB, eventually triggering cytokine release. (Fig. 1A) Hence, it is surprising that these two receptors share the same signal transduction pathway, while having different functions.

**Figure 1.**
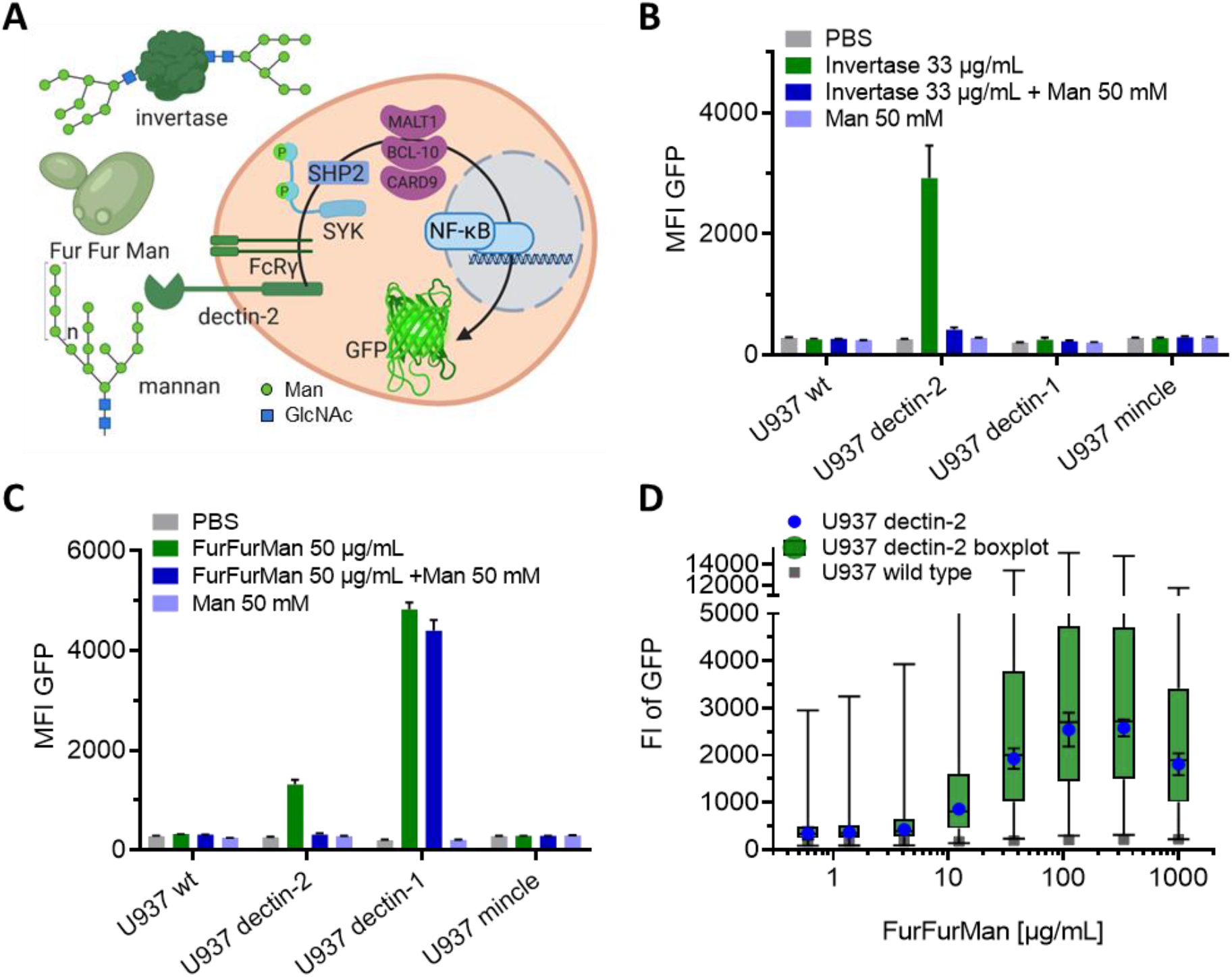
Reporter cell system for observation of glycan-lectin interaction. A) Schematic representation of dectin-2 signalling pathway with GFP under control of NFκB; on the left three ligands for dectin-2 (invertase and mannan from Saccharomyces cerevisiae, and FurFurMan from *Malassezia furfur*). Scheme created with BioRender.com. B) and C) Monoclonal reporter cells expressing dectin-1, mincle, dectin-2, or wild type were stimulated with invertase (B) or FurFurMan (C) n=3. D) Dose response of the dectin-2 reporter cells is shown both as geometric mean with standard deviation and Boxplot with the whiskers representing the 1 percentile of the cellular population (n=6).

In contrast, dectin-1 and dectin-2 have different signal transduction pathways, but are both involved in the detection of fungal infections. They recognize β-glucans and mannan, respectively. Upon fungal infection, combination of these and other cell surface receptors expressed by antigen presenting cells then leads to a defined immune reaction *via* signal integration processes.(Snarr et al., 2017) Such signal integration can result in synergism between the receptors triggering an effect greater that their individual contributions. For example, MCL, another CLR present on cells of the innate immune system and just like dectin-2 an evolutionary offspring of mincle, is known to synergistically work with dectin-2. (Ostrop et al., 2015; Zhu et al., 2013) Additionally, to this type of synergism, other members of the CLR family, *e*.*g*. DC-SIGN and Langerin, rather modulate a response instead of initiating it by themselves.(Geijtenbeek & Gringhuis, 2016; Osorio & Reis e Sousa, 2011) Overall, the resulting complexity of how these cell surface receptors can modulate each other to translate a glycan encoded information into useful cellular signalling it is essential to quantitatively describe their interaction.

Herein we studied DC-SIGN, MCL, dectin-1, dectin-2, mincle, TNFαR and TLR-1&2 in NFκB GFP reporter cells using single cell resolved flow cytometry. To accurately quantify the information these lectins and other receptors transmit we use the channel capacity as metric. The channel capacity is the maximum of information which can be transmitted by an information channel.(Cheong et al., 2011) In our case, an information channel is described by the receptor and its signal transduction pathway which ultimately lead to NFκB translocation and finally GFP expression in the reporter model. Yet the channel capacity isn’t merely describing the resulting maximum intensity of the reporter cells. The channel capacity takes cellular variation and activation across a whole range of incoming stimulus of single cells resolved data into account and quantifies all of that data into a single number. We therefore use the channel capacity to quantify the stochastic system of glycan lectin communication. By employing this metric, we found dectin-2 to stimulate NFκB rather inefficiently due to a wide spread cellular distribution. Furthermore, we found that synergistically acting lectins can enhance channel sensitivity but not the transmitted flow of information. Overall, our findings and approach contribute to a quantitative description of glycan lectin communication and signal integration of CLRs and other receptors, which may lead to a better understanding of key phenomena such as pathogen recognition and autoimmunity.

## Results

### Quantifying signal transduction in glycan-based communication

We employed a single cell resolved reporter system to monitor CLR activity by GFP expression under control of the transcription factor NFκB in human monocytic U937 cells. Dectin-2 was expressed in these reporter cells and stimulation was conducted using various ligands (Fig. 1A). FurFurMan, an extract of *Malassezia furfur*, as well as the polysaccharide mannan and invertase, both from *S. cerevisiae*, initiated dectin-2 signalling. In contrast, owing to the lack of multivalency, mannose itself could not initiate signalling, but was able to inhibit dectin-2 function (Fig. 1B, Fig. 1C, and Supplementary Fig. S1A).(T. Ishikawa et al., 2013) The stimulation of human dectin-2 receptor is in line with previous reports on its murine homolog, which is triggered by Man-α1-2 Man moieties presented on scaffolds like proteins, glycans, or polystyrene beads.(T. Ishikawa et al., 2013; Yonekawa et al., 2014; Zhou et al., 2018). Introduction of dectin-1 into these reporter cells enabled detection of NFκB-based GFP expression. However, while FurFurMan could also stimulate dectin-1 cells, this was not inhibited by addition of mannose, which is expected for this β-glucan receptor (Fig. 1C).

Dose response curves of dectin-2 reporter cells stimulated with FurFurMan revealed that the point of maximal stimulation was not at the highest concentration of ligand (Fig. 1D). Furthermore, the single cell resolution of the assay offered the possibility to study the stochastic behaviour of the population dynamics of the reporter cells: deterministically, the geometric mean suggests a significantly different cellular response for various data points (*e*.*g*. 12.3 vs. 111.1 µg/mL, p<0.001). However, single cell data analysis of the cellular population revealed an overlap with the unstimulated population, even at maximal stimulation exemplifying similar cellular response under altered stimulation. This demonstrates the absence of a clear two-state behaviour on a population level (Fig. 1D, Supplementary Fig. S1B). To rule out any influence of the selection process for the cellular clones, dectin-2 expressing cells were sorted according to their GFP expression level. When re-stimulated, both populations again showed the same broad GFP expression, confirming the wide range of the response to be independent of genetic differences between individual cells (Supplementary Fig. S1D). Taken together, observing dectin-2 signalling on a single cell level in relevant model cell lines reveals a broad population distribution when stimulated.

### Dectin-2 transmits less information than other receptors

To investigate whether other receptors with similar signalling pathways follow the same principle, we analysed the dose response of dectin-1, mincle (another close relative of dectin-2) and the non-CLRs TNFαR and TLR-1 and-2 (Bode et al., 2019; Holbrook et al., 2019; E. Ishikawa et al., 2009; Ozinsky et al., 2000). All receptors but dectin-2 showed an amplitude of the dose response differentiating the maximally stimulated from the baseline population (Fig. 2A, Supplementary Fig. S2A).

**Figure 2.**
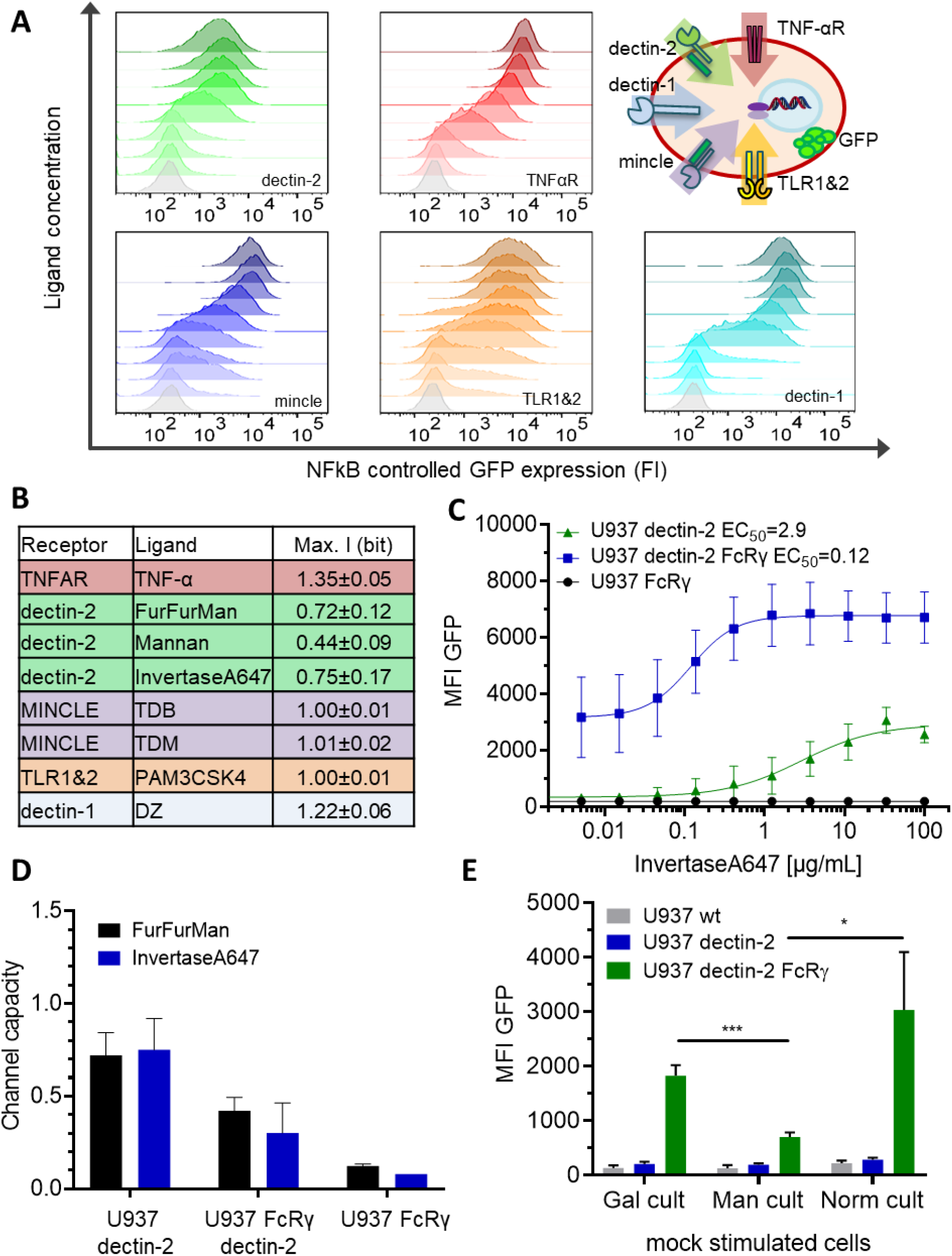
Quantification of signal transduction. A) Representative histograms of U937 reporter cells dose response, stimulated specifically with invertase for dectin-2, TNFα for the TNFαR, TDB for mincle, PAM3CSK4 for TLR1&2, and depleted zymosan for dectin-1. Top right panel shows a schematic representation of the five analysed receptor channels. B) Table summarizing the receptors calculated channel capacities with the respective ligands n≥3 of independent experiments. The channel capacity values given are an average of all experiments we conducted for this study. The respective sample sizes are TNF-α n=8, FurFurMan n=7, Mannan n=5, Invertase647 n=8, TDB and TDM n=3, PAM3CSK4 n=4, DZ n=6. C) Monoclonal reporter cells either expressing dectin-2 (n=3), FcRγ (n=2), or dectin-2 and FcRγ (n=4) were stimulated for 16h with various concentrations of invertase. The 95%CI for the EC_50_ are as follows for dectin-2 1.3-12.3 µg/mL, for dectin-2 FcRγ 0.02-0.27 µg/mL. D) Calculated channel capacities from invertase and FurFurMan stimulation (data also seen in figure C and B) and FurFurMan stimulation of U937 reporter cells. E) Mock stimulated reporter cells 16h after cultivation with 25 mM mannose, or galactose, or under normal conditions for 48h (n=3). Statistical significance determined with an unpaired t-test; with symbols representing ns for p-value > 0.05, * for p-value ≤ 0.05, ** for p-value ≤ 0.01, *** for P ≤ 0.001.

To further characterize and more importantly quantify the underlying signal transmission in a cellular population, the channel capacity was used as a metric. This channel capacity quantifies the maximum amount of information, which can be transmitted via a communication channel. In a biological context the channel is the receptor and its connected signal transduction pathway to the point where the experimental data is recorded. In our system the dectin-2 channel is composed of the receptor dectin-2 and its associated signal transduction pathway all the way to the experimental readout of GFP. (Fig. 1A, Fig. 2A) Previous work on TNF-α signalling found the TNF-α channel to have a channel capacity of about 1 bit in particular 1.64±0.36 bit when a reporter cell system was used.(Cheong et al., 2011) Such a 1 bit channel capacity suggests that a cellular population can use a receptor to distinguish between two states - on/off or presence/absence of a ligand. Based on Cheong *et al*. 2011 (Cheong et al., 2011) we developed a script to calculate the channel capacity from flow cytometry data. We used titration curves of the reporter cells expressing various receptors to quantify the channel capacities of these receptors. (Fig. 2A, Supplementary Fig. S2A, see SI for a detailed description of channel capacity calculations) For U937 cells, we found the TNFαR channel to have the highest capacity of 1.35±0.04 bit, which was not influenced by the introduction of additional lectins (*i*.*e*. mincle, dectin-2, and DC-SIGN, see Supplementary Fig. S2B). Dectin-1 came next at 1.22±0.06 bit, while both mincle and TLR1&2 had a channel capacity of 1.00±0.01 bit. Since all of these receptors signal via NFκB, these differences can be explained by receptor expression levels and downstream pathways. In contrast, dectin-2 stimulation resulted in a channel capacity of 0.72±0.12 bit using FurFurMan as a ligand. Stimulation using heat inactivated invertase or mannan had 0.78±0.15 and 0.44±0.09 bit, respectively (Fig. 2B). Also in THP-1 cells a similar trend of lower GFP expression upon stimulation is observed, further supporting the notion that dectin-2 has a lower transmission efficiency compared to other receptors such as TNFAR (Supplementary Fig. S2C-E). The difference between mincle and dectin-2 is striking as both lectins use the same signalling pathway *via* FcRγ (T. Ishikawa et al., 2013), suggesting that substantial differences between the channel capacities rely on very early ligand recognition events.

We hypothesized overexpression of the signalling protein FcRγ might increase the information transmitted *via* dectin-2. The overexpression of FcRγ resulted in at least twofold increase of this signalling subunit (Fig. 2C). Overexpression of both dectin-2 and FcRγ yielded a high basal NFκB activation of the cells while the channel sensitivity for its ligand (EC_50_) increased about 50-fold (Fig. 2C, Supplementary Fig. S2D). While the maximal signal (MFI) increased from dectin-2 to dectin-2 FcRγ cells, the channel capacity however decreased simultaneously (0.30±0.16 bit) (Fig. 2C, D). From this, we concluded the channel capacity of a glycan-based communication channel is not necessarily coupled to its sensitivity. Also, the ability of a communication channel to transmit information is not well described by its maximal signal alone (*i*.*e*. MFI), but rather by the channel capacity. Next, we quantified the number of receptors and excluded that the difference in mincle and dectin-2 channel capacities are due to differences in receptor expression levels (Supplementary Fig. S2D). Taken together, dectin-2 transmits a relatively low amount of information and while its sensitivity (EC_50_) can be modulated with FcRγ, the transmitted information does not increase. Additionally, the number of receptors has little influence on the channel capacity or amplitude.

### Signal integration compromises between lectin receptors when both are engaged

To expand our insight from isolated cell surface receptors to the interplay between multiple lectins we prepared reporter cells expressing dectin-2 and dectin-1 simultaneously FurFurMan served as a stimulant since it interacts with both dectin-1 and dectin-2. First of all, we found that the level of receptor expression did not change upon expression of an additional lectin (Fig. 3A). Dectin-1 expressing cells gave a higher maximal signal (*i*.*e*. maximal MFI) and channel capacity than dectin-2 expressing cells, however the latter channel showed higher sensitivity (EC_50_) to FurFurMan. We found that the double positive cells did compromise between the two receptors displaying the lower EC_50_ of dectin-2 as well as the higher channel capacity of dectin-1 (Fig. 3B, C). Additionally, mannose could be used to interfere with dectin-2 signalling, thus U937 dectin-1 dectin-2 expressing cells showed the same dose-response curve as dectin-1 expressing cells (Fig. 3D). When depleted zymosan (DZ), a dectin-1 specific ligand, was used, dectin-2 expression did not significantly influence the response of the double positive cells. Hence, dectin-2 specific signalling was not influenced by dectin-1 expression (Supplementary Fig. S3A-C). From these two lectins we see that multiple receptors stimulated simultaneously resulted in a compromise between their channels, which demonstrates the importance of cross talk between lectins.

**Figure 3.**
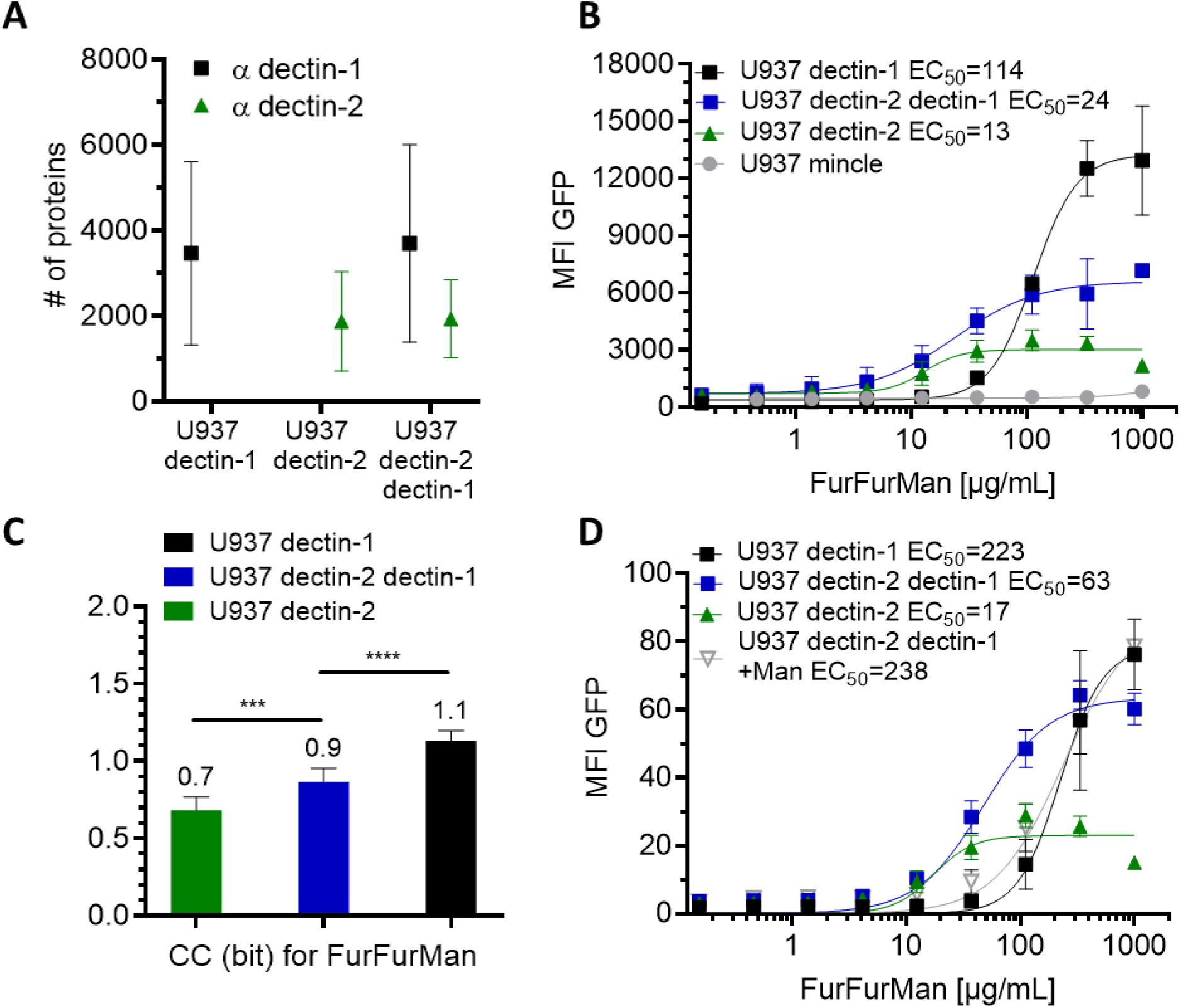
Signal integration of dectin-1 and dectin-2. A) Quantitation of surface expression of U937 dectin-1, dectin-2, and dectin-1 dectin-2 U937 reporter cells. FI values were transformed into the number of proteins expressed using a PE-quantitation kit. Graph shows geometric mean ± robust SD of the cellular population. B) Monoclonal reporter cells either expressing mincle, dectin-2, dectin-1, or both dectin-2 and dectin-1, (n≥3) were stimulated for 16h with various concentrations of FurFurMan. The 95%CI (profile likelihood) for the EC50 are as follows for dectin-1 99-132 µg/mL, for dectin-2 9-18 µg/mL, for dectin-2 dectin-1 16-42 µg/mL. C) Channel capacities of U937 reporter cells expressing either dectin-1, dectin-2, or both stimulated with FurFurMan. Statistical significance determined with an unpaired t-test, n=9. D) Monoclonal reporter cells either expressing dectin-2, dectin-1, or both dectin-2 and dectin-1, (n=3) stimulated with various concentrations of FurFurMan. Cells were stimulated either with or without 25 mM mannose, shown are mean ± SD of n=3.

### Synergistic receptors cannot enhance the information transmitted by dectin-2

To relate the amount of incoming stimuli with the cellular response, we included binding data in our channel capacity calculation. We applied fluorescently labelled ligands, TNF-α and invertase, which enabled us to directly monitor cellular conversion of labelled ligand stimulus to output. Thus rather than having discrete concentration steps, we had a continuous readout for the ligand. We ensured that dye conjugation did not alter the channel capacity and found the channel capacity calculated from a discrete titration curve or the continuous directly labelled ligand to be the same (Supplementary Fig. S4A, S4B). Although this could simply be a result of delayed GFP expression, we noted that for both TNFαR and dectin-2, the cellular response saturated prior to receptor saturation (Fig. 4A). We also found that while dectin-2 cells specifically reacted to invertase, they did not bind more invertase than wt cells (Supplementary Fig. S4C, D). Yet even when directly monitoring cellular conversion of labelled ligands to output, dectin-2 channel capacity stayed low relative to TNF-αR (Supplementary Fig. S4B).

**Figure 4.**
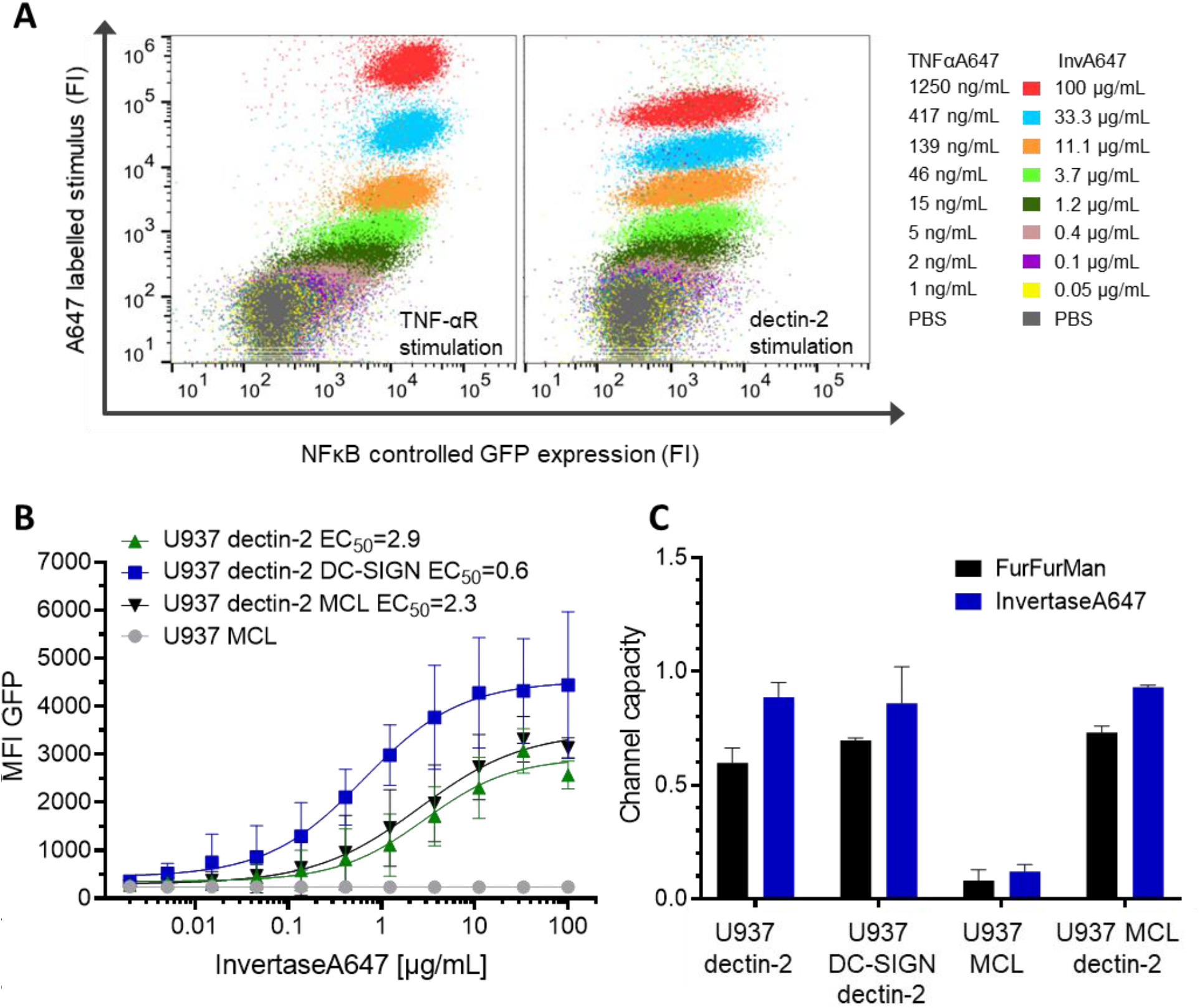
Effects of ligand labelling and synergistic lectins on dectin-2. A) 2D plots of single cells resolved dose responses of U937 dectin-2 reporter cells. Left: cells were stimulated with Atto647 labelled TNF-α. Right: cells were stimulated with Atto647 labelled invertase. Colours from red to grey represent titration of the ligands. B) Dose response of Atto647 labelled invertase stimulation of U937 cells expressing dectin-2 alone, in combination with DC-SIGN, or in combination with MCL n≥3. The 95%CI (profile likelihood) for the EC_50_ are as follows for dectin-2 1.3-12.3 µg/mL, for dectin-2 DC-SIGN 0.2-1.9 µg/mL, for dectin-2 MCL 1.0-16.8 µg/mL. The EC_50_ of dectin-2 DC-SIGN did significantly differ from dectin-2 (p=0.0415), but dectin-2 MCL did not. C) Channel capacities of dectin-2 in combination with DC-SIGN and MCL after stimulation with either FurFurMan or invertase n≥3.

We wondered whether dectin-2 needs a partner to synergistically increase its signalling, so we included DC-SIGN and MCL (Fig. 4B). Although DC-SIGN does not elicit signalling by itself in U937 cells, it is known to recognize high mannose structures present on invertase.(Gringhuis et al., 2009) As expected, U937 dectin-2 DC-SIGN cells experience significantly increased ligand binding (Supplementary Fig. S4C, D). We then speculated that this would either a) aid the recognition by pre-concentration of the ligand on the plasma membrane or b) would serve as a decoy receptor decreasing stimulation of dectin-2. In fact, DC-SIGN mediated ligand binding did not alter the dectin-2 channel capacity for FurFurMan or invertase stimulation nor did DC-SIGN expression itself modulate TLR4 signalling (Fig. 4C, Fig. S4F). Additionally, we investigated overexpressed MCL, a CLR known to synergistically promote dectin-2 signaling.(Zhu et al., 2013) MCL expressing cells did not respond or bind more invertase and no synergistic effect with dectin-2 was observed (Fig. 4B, C Supplementary Fig. S4D). We then wondered whether the difference in channel capacity between dectin-2 and TNFαR could simply be a result of affinity. Since TNFαR has a nanomolar affinity for its ligand (Grell et al., 1998), we applied an anti-dectin-2 antibody to stimulate dectin-2 cells. Even under these conditions, we did not monitor an increase in channel capacity beyond the 0.74 bit (Supplementary Fig. S4E). Therefore, we found that neither DC-SIGN nor MCL significantly increase dectin-2 channel capacity, but DC-SIGN enhances its sensitivity and cellular binding of invertase.

### Dectin-2 signalling is accompanied by inherently high noise

A striking feature throughout this study that distinguishes the high capacity TNFαR channel from dectin-2, is the broad distribution of the response of dectin-2 cells upon stimulation (Fig. 5A). A commonly used measure for such variability of a cell population, or rather cellular noise, is the squared normalized standard deviation (CV^2^).(Colman-Lerner et al., 2005) We therefore used the data obtained from the dose response measurements and quantified the noise. While TNFαR and mincle showed low noise at their point of maximal stimulation, dectin-2 noise increased and remained at elevated levels during titration of both FurFurMan and invertase (Fig. 5A, Supplementary Fig. S5A). In addition, the high channel noise of TLR1&2 and the low noise of dectin-1 were able to reach a relative minimum at maximal stimulation (Supplementary Fig. S5A). To exclude cell-to-cell variation in protein expression we subsequently compared the GFP response upon stimulation with the constitutively expressed mAmetrine, which originally served as a marker for vector presence in the cells. We found that NFκB activation did correlate with mAmetrine while other parameters like cell morphology (FSC and SSC) or cell cycle did not (Fig. 5B and Fig. 5C). Interestingly, the cellular response of TNFαR and mincle did correlate with mAmetrine (R^2^ of 0.68 and 0.67), in contrast to dectin-2 (R^2^ of 0.31, Supplementary Fig. S5C). We accounted the NFκB triggered GFP expression noise for the expression noise of the unrelated constitutively fluorescent protein mAmetrine (Colman-Lerner et al., 2005). Nevertheless, the noise level of the dectin-2 channel stayed significantly higher than that of TNF-α and mincle (Fig. 5C and D). Thus, dectin-2 NFκB signalling is accompanied by high noise, but not as a result of protein expression noise, cell morphology, or cell cycle. This high noise stops dectin-2 from transmitting more information in the cellular population.

**Figure 5.**
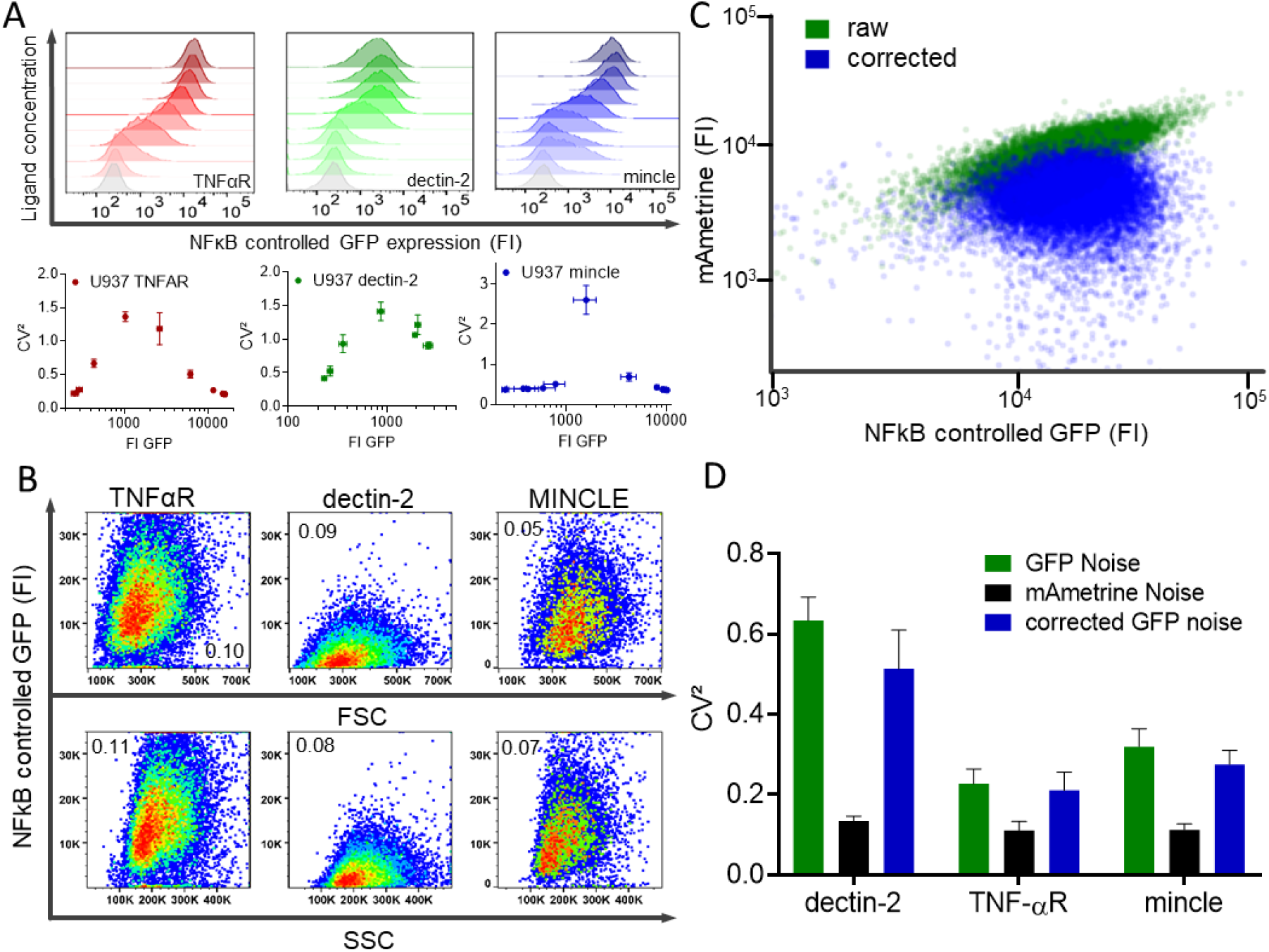
Noise quantification of signalling receptors. A) Top row shows histograms seen in Fig. 2 A. Bottom row shows noise quantification of U937 reporter cells dose responses either by TNF-αR, dectin-2, or mincle stimulation. Y-axis shows CV^2^ (Robust standard deviation/ geometric mean)^2^ to quantify noise of the cellular population. The x-axis shows fluorescent intensity of the NFκB expressed GFP. The last three data points were compared using an unpaired t-test and for all three cases dectin-2 was significantly different from TNF-α and mincle p-value ≤ 0.01. B) 2D single resolution plot of constitutively expressed mAmetrine and NFκB controlled GFP. Plotted are raw data and data corrected for mAmetrine correlation, which was achieved by turning every raw data point by the angle of a linear correlation to the x-axis. This demonstrates how we corrected for general protein expression noise. C) Density plots of NFκB controlled GFP and either forward scatter (FSC) or mAmetrine. Number gives R^2^ for a linear regression of each plot. D) Noise quantification of U937 reporter cells by TNF-αR, dectin-2, or mincle stimulation, cellular noise was quantified using the CV^2^ (standard deviation/mean)^2^ of the cellular population either for GFP, mAmetrine, or GFP corrected for protein expression noise as demonstrated in C).

## Conclusion

We set out to better understand how glycan-encoded information is read in cell-cell communication. We established an *in vitro* model system and applied the channel capacity as a quantitative metric. For the receptors other than dectin-2 it was not surprising to find channel capacities of around 1 bit, since similar values have been reported for other systems previously (Suderman et al., 2017). In particular, TNF-α receptor has a channel capacity of 1.64±0.36 bit, which was found in a comparable reporter cell system.(Cheong et al., 2011) Interestingly, the number of receptors expressed on the cell surface did not determine the channel capacity of a signalling channel (Supplementary Fig. S2D). Our results exemplify that lectin signalling pathways and especially the dectin-2 pathway should not be viewed as a deterministic on/off-switch, but rather as difference in the probability of cells to be active at a certain dose (Fig. 1D). This is in line with previous reports strengthening a quantitative view of cellular signalling and taking the cellular microheterogeneity into account.(Levchenko & Nemenman, 2014; Zhang et al., 2017) We found that the mannose binding CLR dectin-2 to transmits less information compared to other receptors of the same family or even unrelated immune cell receptors. (Fig. 2A&B). (Cheong et al., 2011; Suderman et al., 2017)

To understand how these insights could be expanded on the interplay between multiple receptors like the CLRs occur on innate immune cells rather than isolated lectins, we employed combinations of CLRs on our model cells. Dectin-2 and dectin-1, recognize different epitopes on FurFurMan and we found that the effects were not additive, but a compromise between the two receptors, showing the low EC_50_ of dectin-2 but with a higher channel capacity derived from dectin-1. Next, the channel capacity of dectin-1+ dectin-2+ cells was a neat average of the individual channel capacities (Fig. 3A&B). This effect is striking since it implies at high concentrations of FurFurMan the dectin-2 channel is actively inhibiting dectin-1 signalling, resulting in a lower cellular NFκB activation. In further experiments, we showed that dectin-1 and -2 themselves are not influencing one another when stimulated with specific ligands (Fig. 3C&D). This demonstrates that when multiple lectins with shared pathways are engaged simultaneously, the cells will integrate both channels by compromising between them rather that favour one over the other or combining the two channels. It is well known that lectins are able to modulate the signals of other receptors.(Geijtenbeek & Gringhuis, 2009; Gringhuis et al., 2009; Miyake et al., 2015) Yet this compromise is an exciting discovery since to the best of our knowledge previous studies have not quantified lectin signal integration. Hence, it is likely that during a fungal infection, exposing multiple epitopes, the precise arsenal of immune receptors and their underlying pathways are integrating the information contained within the carbohydrate and non-carbohydrate ligands. This in turn leads to a compromise of all activated receptors and results in a specifically tailored biochemical response of the given immune cell.(Ostrop & Lang, 2017)

Dectin-2 itself we found to be relatively inefficient when compared to the closely related mincle that uses the same pathway more effectively. (Fig. 2A&B) It is therefore likely the receptor itself determines very early on the information flow into the cell. This could be a result of mincle being stimulated with crystalline insoluble ligands which could result in larger signalling clusters at the cellular surface. Alternatively, dectin-2 signalling could be influenced by mannose structures which are present on the cellular surface by that giving rise to background signalling and selection for inefficient signalling in an in vitro setting of high cellular density (Supplementary Fig. S2E). Additionally, since dectin-2 binds high mannose structures of eukaryotic origin,(McGreal et al., 2006) a too sensitive reaction might lead to permanent self-recognition of human Man9 structures for example and hence potential autoimmune reactions. This hypothesis is supported by the dectin-2-dependent high basal activity of dectin-2 FcRγ cells, which in turn is responsible for a lower channel capacity in dectin-2 FcRγ cells (Fig. 2C-E). Hence, dectin-2 could have evolved to use the CARD9-BCL-10-Malt1 pathway to NFκB less effective. Along the same lines recent reports show that CLRs are in general becoming more important in autoimmunity, dectin-2 in particular is known to be responsible for the development of allergic reactions.(Dambuza & Brown, 2015; Parsons et al., 2014) To incorporate binding of the stimulus into the channel capacity calculation, we labelled two stimulants. We observed both ligand and cellular reaction on a single cell level simultaneously (Fig. 4A). Interestingly, dectin-2 did not seem to bind more ligand, but was responsible for NFκB activation. We first thought a combination of multiple lectins might synergistically enhance signalling capacity of dectin-2. But while DC-SIGN greatly enhanced ligand binding to the cells, it did not significantly increase in channel capacity. However, the presence of a second mannose-binding receptor increased the channels sensitivity (EC_50_) (Fig. 4B&C, Supplementary Fig. S4C). We also found that in contrast to murine dectin-2, the closely related lectin MCL (Dectin-3) did not have a significant synergetic effect on human dectin-2 signalling. Based on those results we hypothesize that channel capacity of a lectin is predetermined by the receptor itself. In contrast, its channel sensitivity (EC_50_) can be enhanced by synergistic receptors or signalling molecules. The maximally transduced information cannot (Fig. 2C&D and Fig. 4B&C).

To gain deeper insight into the inefficiency of dectin-2 mediated cell activation we quantified and investigated the origin of its high noise and saw that it was not related to cell morphology (FSC, SSC) or cell cycle (Fig. 5B, Supplementary Fig. S5B&C). Moreover, dectin-2 related noise stayed high even after accounting for expression noise (Fig. 5D).(Colman-Lerner et al., 2005) We conclude that dectin-2 by itself is inherently noisy and an inefficient transmitter of information. The reason for this might simply be as mentioned before a more effective dectin-2 could result in autoimmune defects. It would be exciting to see a quantitative characterization of receptor channels in primary cells, since this could reveal how combination and integration of various channel leads to distinct cellular responses in the immune system. Since channel capacity calculations are applicable regardless of the nature of signal and medium, (Levchenko & Nemenman, 2014) one could use it to objectively quantify cellular responses in similar assays in the future.. This will help to better characterize communication channels especially in such a complex and enigmatic field of study as glycan lectin interactions.

## Materials and Methods

All reagents were bought from Sigma Aldrich, if not stated otherwise.

### Reporter cell generation and reporter cell assay

U937 cells (a kind gift from Dr. Nina van Sorge, UMC Utrecht, the Netherlands) were transduced with an NFκB-GFP Cignal lentivirus (qiagen) according to the manufacturer’s instructions to generate NFκB reporter cells. 0.5 mL of 2e5 cells were mixed with the lentivirus at an MOI of 15 and spin transduced for 1.5 h at 33°C and 900g. After 48h of rest, cells were selected with puromycin (gibco) for 3 passages. Eight cultures from a single cell each were subsequently made and evaluated according to their GFP expression, clone #5 was chosen and used for all experiments of this paper.

### Reporter cell assay

U937 reporter cells were used in its log phase and 100µL were plated in a 96-well plate with 3e4 cells per well. Cells were challenged in complete media (RPMI with 10% FBS, 1% Glutamax, 1% Pen/Strep, all by gibco) with TNF-a and various other ligands and at various concentrations for 16h and 13h, respectively. After incubation, cells were re-suspended once in DPBS and the expressed GFPs fluorescent intensity was measured by flow cytometry (Attune Nxt, Thermo Fisher).

### Cell culturing and passage

U937 cells were kept between 1e5 and 1.5e6 cells/mL in complete media with passage 2-4 times a week. 293F cells were adherently cultured in DMEM with 10% FBS, 1% Glutamax, 1% Pen/Strep, (all gibco) and split 2-3 times per week. All cells were tested for mycoplasma contamination using Minerva biolabs Venor®GeM Classic.

### Generation of lectin overexpressing cells

cDNA of MINCLE, dectin-2, MCL, FcRγ, dectin-1, and DC-SIGN were cloned into vector BIC-PGK-Zeo-T2a-mAmetrine:EF1a as previously reported.(Wamhoff et al., 2019) This bicistronic vector expresses mAmetrine under the PGK promoter. To combine multiple GOI we also used the lentiviral vector EF1a-Hygro/Neo a gift from Tobias Meyer (Addgene plasmid # 85134). Briefly, 293F cells were transfected with vectors coding for the lentivirus and GOI. Lentivirions were generated for 72h and the supernatant was frozen to kill any remaining 293F cells. This supernatant was used to transduce the GOI into U937 cells *via* spin infection at 900g and 33°C in the presence of 0.8µg/mL polybrene(van de Weijer et al., 2014). After 48h rest, the U937 cells were selected with appropriate antibiotics (Zeocin 200µg/mL, G418 500 µg/mL, or Hygromycin B 200µg/mL; Thermo Fisher, Carl Roth, Thermo Fisher, respectively). A list of used primers can be found in the Supplementary information.

### Antibody staining and quantitation

For the surface staining, cells were incubated in with the respective antibodies and isotype controls for 30 minutes at 4°C in DPBS, then washed once in DPBS +0.5% BSA and measured via flow cytometry. For perforated stains cells were first fixed in 4% PFA (Carl Roth) at 4°C for 20 minutes, then perforated in perforation solution (DPBS + 0.5% BSA + 0.1% Saponine) for 20 min at 4°C. The cells were then re-suspended in perforation solution containing the respective antibodies, incubated for 20 minutes at 4°C, and measured via flow cytometry after being washed once. To quantify the fluorescent intensities, we used the BD PE quantitation kit, which allowed us to calibrate FI to the number of PE molecules present in a sample. A list of all used antibodies can be found in the Supplementary information.

### Labelling of Proteins

Invertase (5 mg in 1 mL) was heat inactivated for 40 min at 80°C and mixed with 3x molar excess of Atto647N-NHS dye (AttoTech) according to the manufacturer’s protocol. The labelled protein was purified using Sephadex G-25 column and aliquots were frozen at -80°C. Since we found the labelled invertase to contain less impurities (Supplementary Fig. S4A) we used Atto647 labelled invertase for all experiments shown in this study. Human TNF-α (Peprotech) was labelled with the same procedure, yet without heat inactivation. The degree of labelling was determined to be around 1 as determined with a labelled protein concentration measurement of a NanoPhotometer NP80 (Implen).

### Channel capacity calculation

Calculations of channel capacity were based on Cheong *et al*. 2011(Cheong et al., 2011) with the only major difference being, that we employed logarithmic binning rather than linear, since we found this to be more fitting to the data.

To calculate the channel capacity from discrete, finite input and output data set, the data set should be projected onto two dimensional, finite rectangular grid plane where the marginal and joint probability can be calculated by counting the number of data points in the individual rectangles. In the dose response of cells, the doses are predetermined and thereby the marginal probability of inputs (i.e., doses) is written as

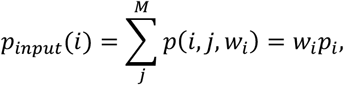

where *p*_*i*_ is the marginal probability of *i*^th^ input, *j* is the index of output, *w*_*i*_ is the weighting value of the *i*^th^ input and *M* is the number of output binning. Therefore, the input entropy can be written as follows:

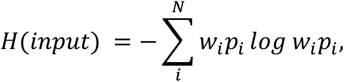

where *N* is the number of input binning. Likewise, the marginal probability of output and corresponding entropy are as follow, respectively:

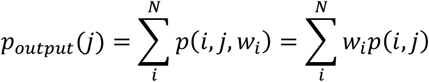

and

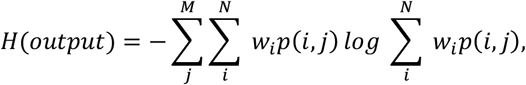

where *p*(*i, j*) is the joint probability of *i*^h^ (input) and *j*^th^ (output) index. Finally, from the joint entropy

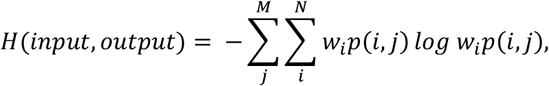

mutual information is calculated as follows:

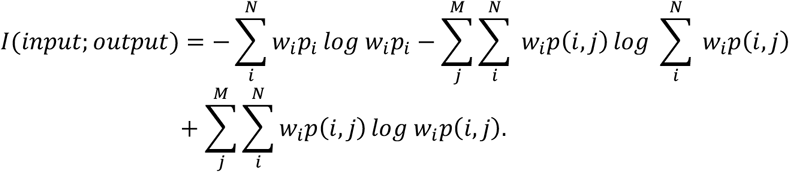

Now the mutual information for the given input and output is the function of *w*_*i*_, and the channel capacity is the maximum value of the function. With the normalization condition 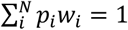, as the constraint, several maximization algorithms can find the input distribution *w*_*i*_ that maximizes the mutual information. We used scipy.optimize.minimize from Python for the maximization.

Supplementary Information Fig. S6 A) and B) show the calculated channel capacity from TNF-α (blue) and dectin-2 (green) channels under the logarithmic A) and linear B) output binning method. Prior to the calculation, the 2% of front-end and 2% tail-end of output data sets were excluded. In the case of logarithmic binning, the channel capacity reaches its plateau from around 2^3^ to 2^9^ binning numbers. Linear binning also gives a stable channel capacity throughout binning number but less value than the logarithmic binning, is due to the logarithmic response of outputs. The relatively wide range of binning numbers which wind up in the similar channel capacity is due to the large sample size (∼150,000). This implies that the sample size is adequate to produce the reliable channel capacity. Supplementary Information Fig. 6 c) also validate the adequacy of sample size by which the random subsampled data sets are calculated into the similar channel capacity value for both dectin-2 and TNF-α.

## Data representation, software, and statistical analysis

Data is shown as mean ± standard deviation. Statistical analysis of data was performed by unpaired two-tailed t-test, with significant different defined as (P < 0.05). EC_50_ values were calculated in graph pad prism version 8.4.2 using four parametric dose vs. response function. When necessary statistical differences between EC_50_ values were compared using an extra-sum-of-squares F test. Detail of statistical tests and EC_50_ determinations can be found in the SI raw data file. FlowJo v.10 was used for analysis and export of flow cytometry data.

## Data Availability

Flow cytometry raw data is available on Dryad. All scripts used and example data are available at: https://github.com/FFuchsbe/GlycanComm

## Acknowledgments

This project (GLYCONOISE) has received funding from the European Research Council (ERC) under the European Union’s Horizon 2020 research and innovation programme (Grant agreement No. 716024). We thank Max Planck Society for support, Prof. Dr. Peter H. Seeberger and Rob van Dalen for helpful discussions. We also thank the Deutsches Rheuma Forschungszentrum (DRFZ) for providing access to their cell sorting facility.

---

**Figure S1.**
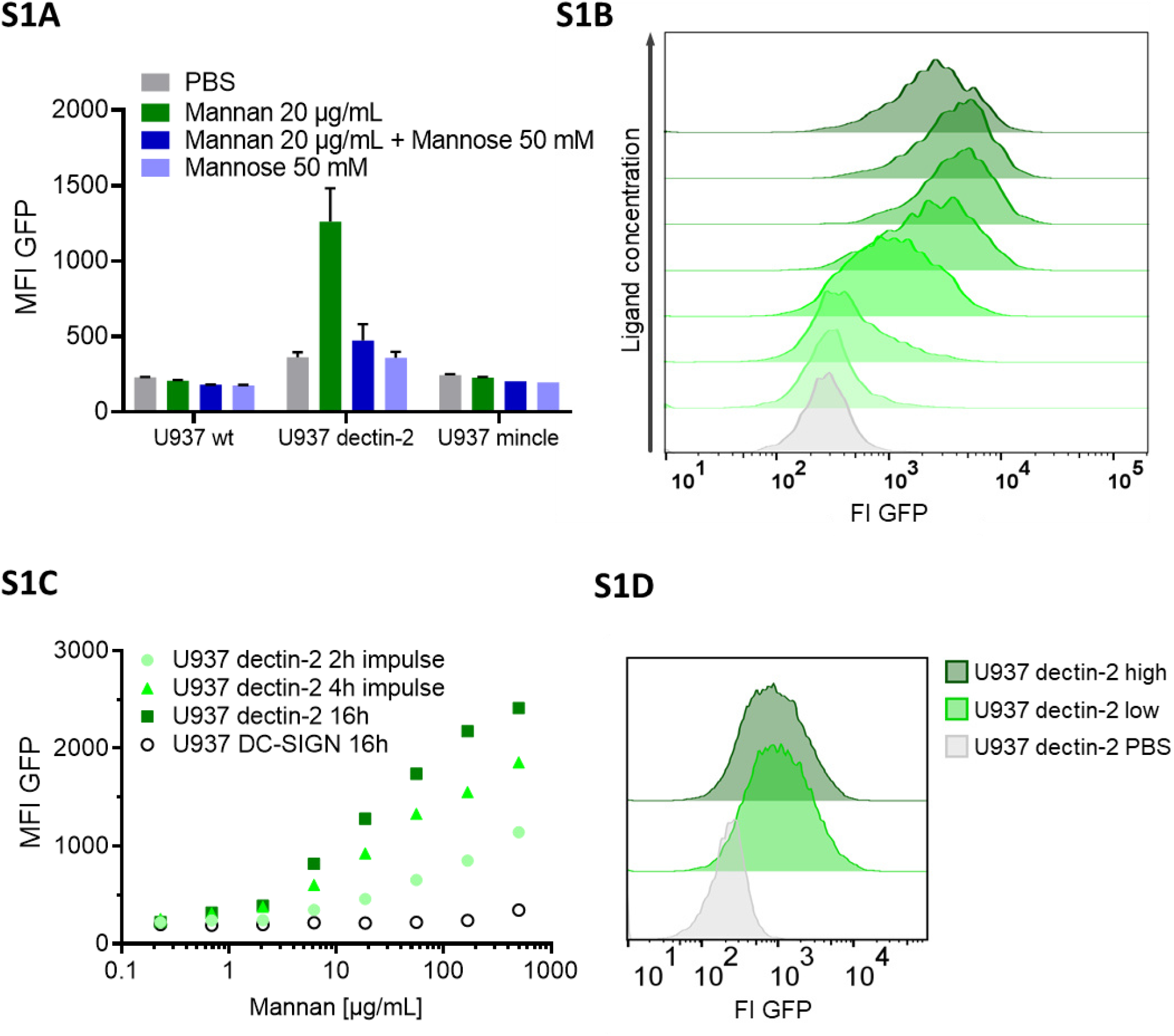
S1A) Monoclonal U937 reporter cells expressing mincle, dectin-2, or wild type were stimulated with mannan (n=4). Mannose alone could not stimulate decin-2, but could inhibit stimulation by Mannan. S1B) Histograms of the dose response in figure 1D, U937 dectin-2 expressing reporter cells react to various concentrations of FurFurMan. Darker histograms were stimulated with higher ligand concentration.S1C) Dose response of dectin-2 and DC-SIGN expressing reporter cells stimulated for 16h, or stimulated for 2 and 4 h, washed in fresh media and incubated to a total of 16h. S1D) U937 dectin-2 reporter cells were sorted in a GFP high and low population after stimulation for 16h with 300 µg/mL Mannan. The sorted cells were the re-stimulated two weeks later with 500 µg/mL.

**Figure S2.**
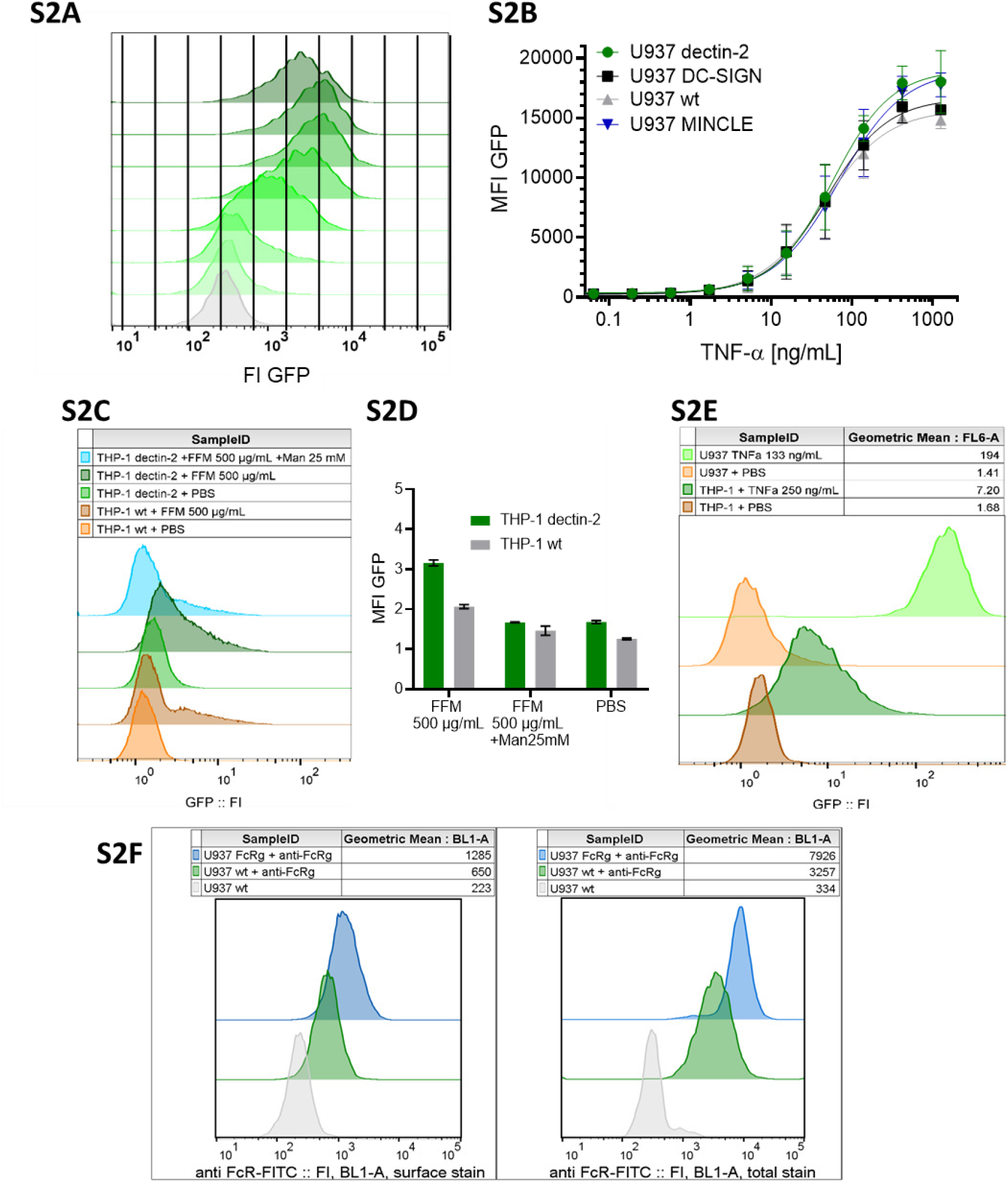
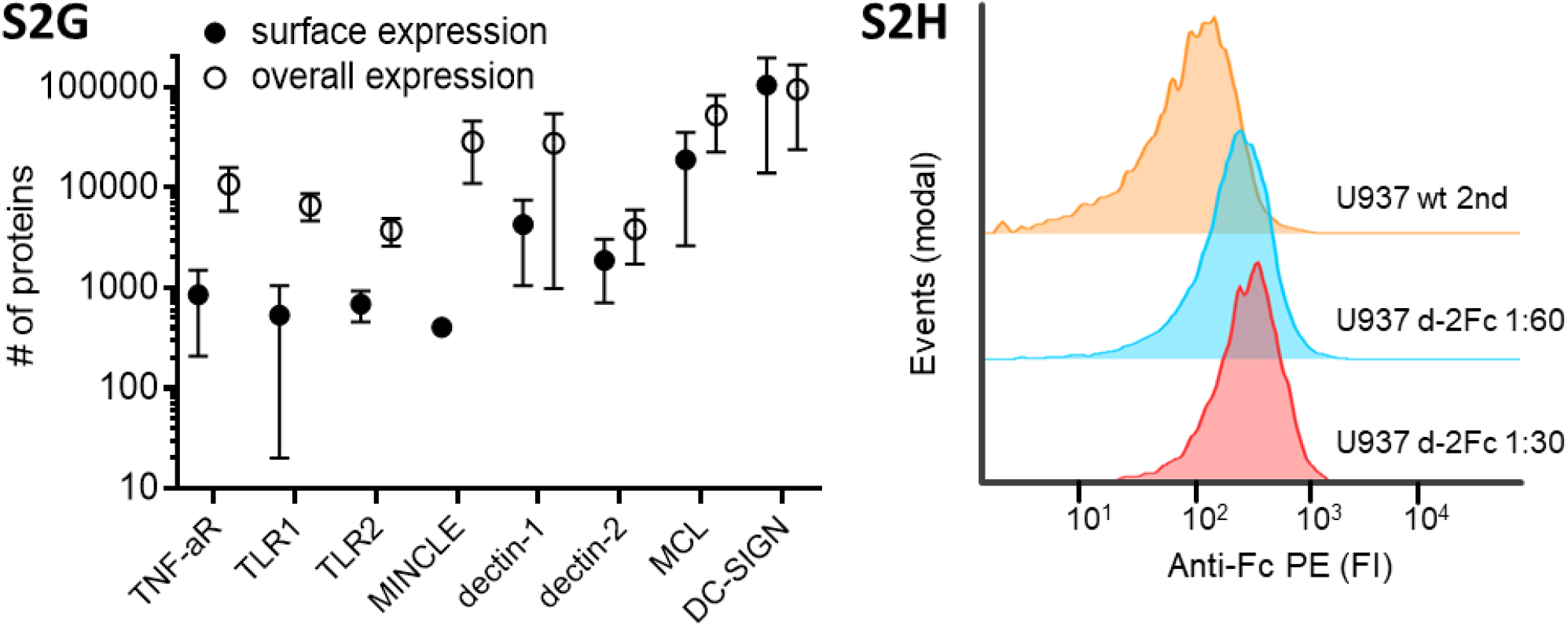
S2A) Histograms of the dose response seen in Figure 1B, with lines demonstrating the binning procedure for calculating the channel capacity. The script “NFKB response_discrete_input_3.py” will use the histograms and maximize the mutual information between input dose and cellular response. While the input is already binned via the number of samples, the cellular response is binned into discrete logarithmic bins. As the dose of ligand increase, the cells have a higher probability of occupying bins of higher cellular reaction or the response increases with dose. S2B) Reporter cell lines expressing various lectins stimulated with TNF-α. S2C) THP-1 reporter cells expressing dectin-2 or wt were stimulated for 48h with FurFurMan (FFM), unstimulated (PBS), or the FurFurMan stimulation was inhibited with 25 mM mannose. Graph shown representative histograms S2D) Geometric means of the experiment done in A in triplicates (n=3) with the error bar representing standard deviation. S2E) Representative histograms showing the TNF-a stimulation (16h) of U937 and THP-1 reporter cells. THP-1 cells stimulated for 48h with TNF-a gave less signal than at 16h (data not shown). S2G) Quantitation of surface and overall expression of receptors used in this study in U937 reporter cells. Cells were stained either for their surface expression or their overall protein expression with PE coupled antibodies. FI values were transformed into the number of proteins expressed using a PE-quantitation kit. Graph shows geometric mean ± robust SD of the cellular population. S2H) Histograms of an anti-FcRγ stain left surface stain, right overall protein levels with a total stain.

**Figure S3.**
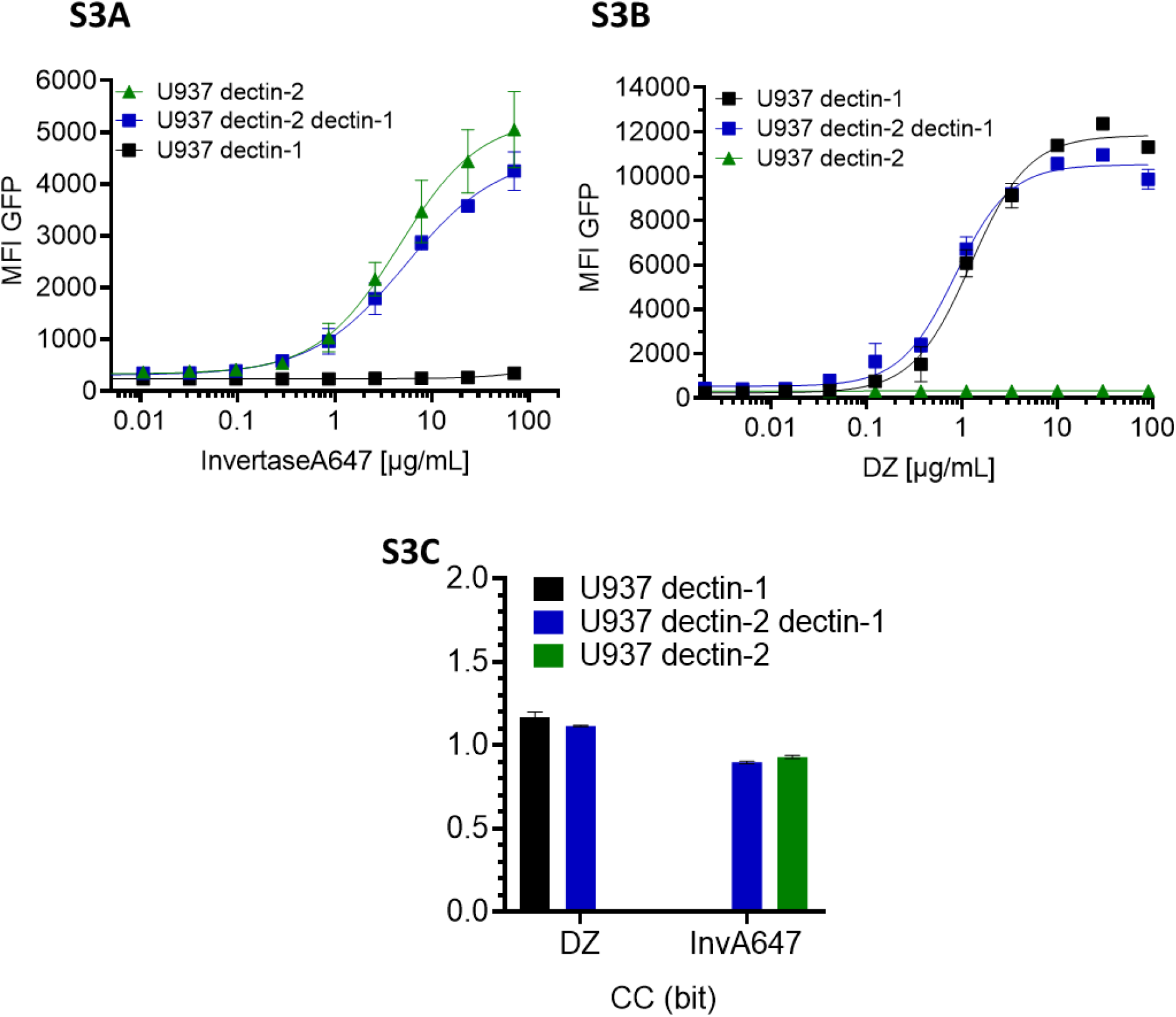
Monoclonal reporter cells either expressing dectin-2, dectin-1, or both dectin-2 and dectin-1, (n≥3) were stimulated for 16h with various concentrations of S3A) InvertaseA647, or S3B) depleted zymosan respectively. S3C) Channel capacities of the data in S3A) and S3B).

**Figure S4.**
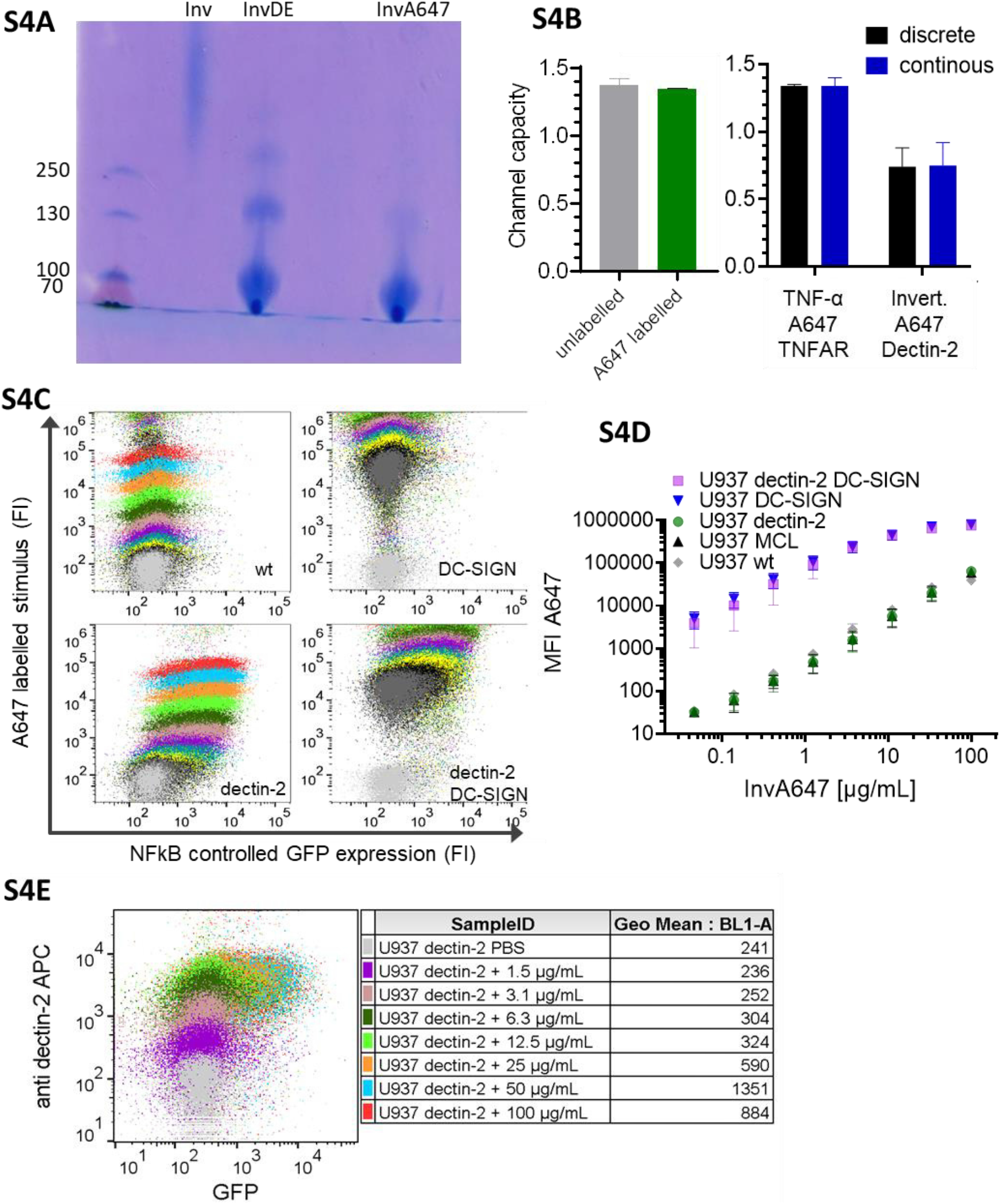
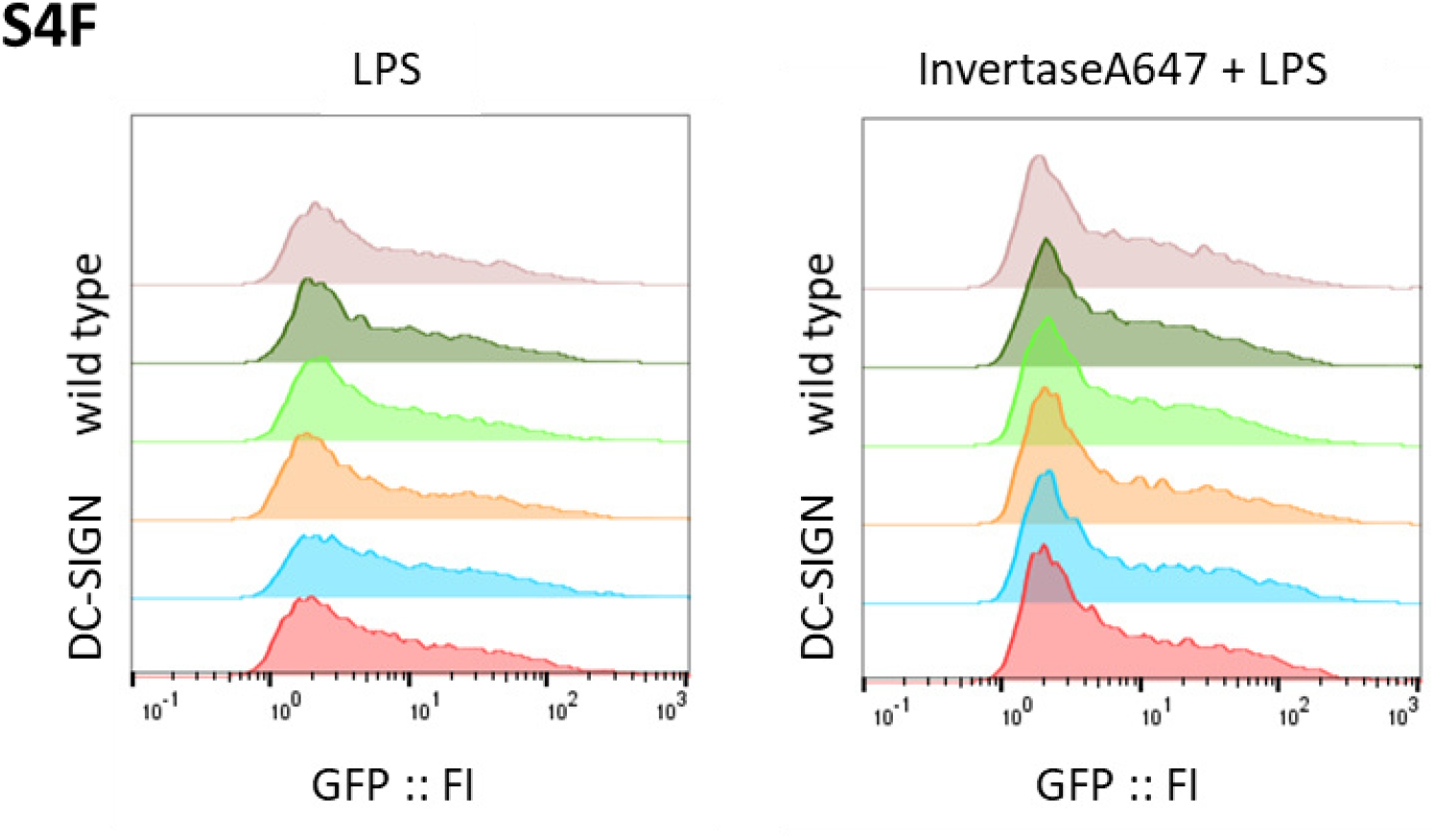
S4A) SDS gel of invertase either native (Inv), or denaturated for 40 min at 80°C (InvDE), or denaturated as before and the labelled with Atto647 (InvA647). Invertase consists of multiple subunits of 135 kDa and 60 kDa.(2) S4B) Left: A647 labelled and unlabeled TNF-α result in the same channel capacity in U937 reporter cells. Right: Channel capacities calculated from discrete or continuous (labelelled) dose-response curves are the same. S4C) U937 reporter cells expressing lectins as indicated, representative 2D plots of dose responses seen in figure 3B. S4D) Comparison of channel capacities for Atto647N labelled Invertase or TNF-α, calculated with discrete titrations of ligand or with using the discrete A647 FI instead. n≥5 bars represent mean ± SD. S4E) 2D dose response of dectin-2 U937 reporter cells stimulated with anti dectin-2 for 16h. S4F) U937 reporter cells either wild type or DC-SIGN expressing were stimulated with 5 µg/mL LPS-EB (invivogen) and 50 µg/mL InvertaseA647 for 18h.

**Figure S5:**
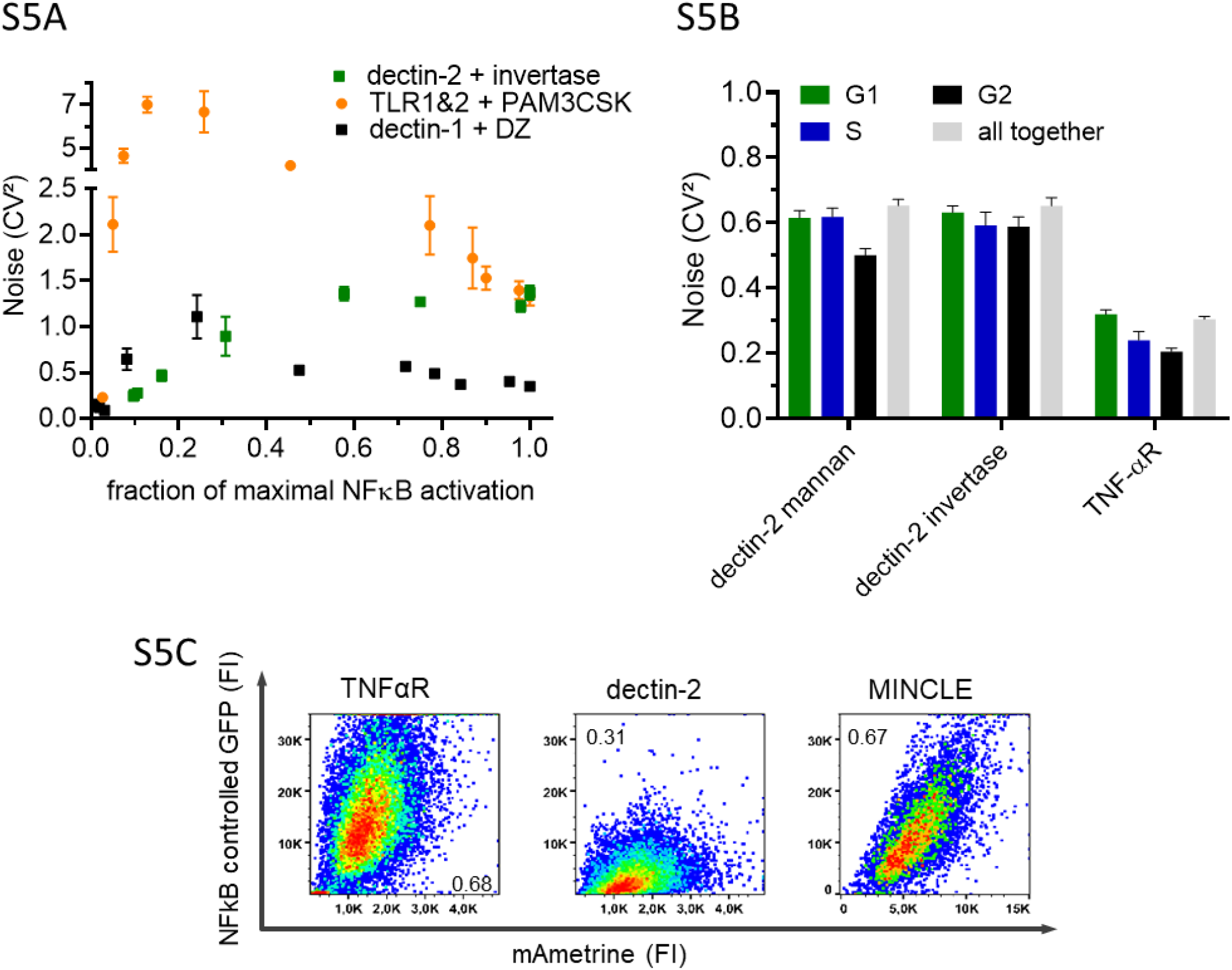
S5A) Noise quantification of U937 reporter cells dose responses either by dectin-2 stimulation with invertase, TLR1&2 stimulation with PAM3CSK4, dectin-1 stimulation with depleted zymosan (DZ) all n≥3. Y-axis shows CV^2^ (Robust standard deviation/ geometric mean)^2^ to quantify noise of the cellular population. The x-axis shows the fraction of maximal stimulation relative to the maximal stimulation of the channels. S4B) U937 dectin-2 expressing reporter cells were stimulated with TNF-α or invertase and gated according to their DNA content and therefore mitotic state (n=3). S4C) Density plots of NFκB controlled GFP and mAmetrine. Number gives R^2^ for a linear regression of each plot.

**Figure S6:**
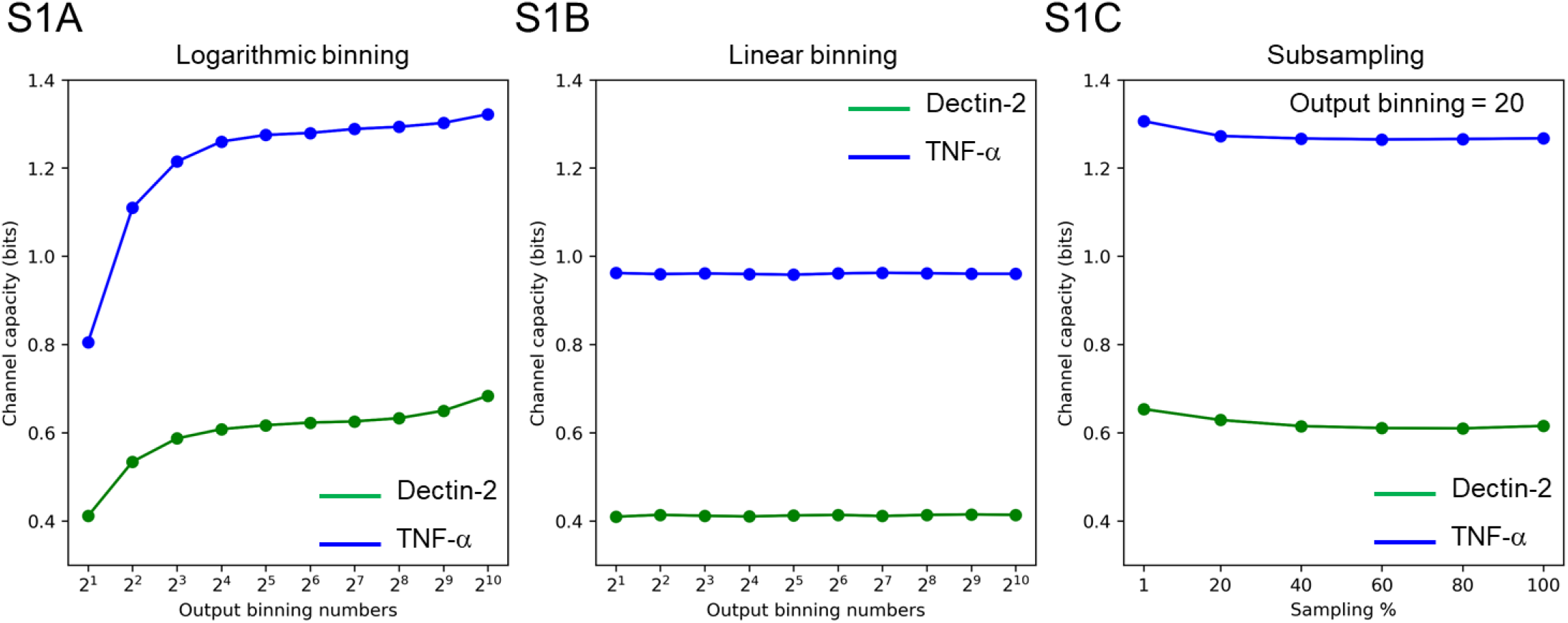
Calculated channel capacity from logarithmic (S6A) and linear binning (S6B) and subsampling (S6C) both for dectin-2 (green) and TNF-α (blue).

